# Dissecting tumor cell programs through group biology estimation in clinical single-cell transcriptomics

**DOI:** 10.1101/2021.10.22.465130

**Authors:** Shreya Johri, Kevin Bi, Breanna M. Titchen, Jingxin Fu, Jake Conway, Jett P. Crowdis, Natalie I. Volkes, Zenghua Fan, Lawrence Fong, Jihye Park, David Liu, Meng Xiao He, Eliezer M. Van Allen

**Affiliations:** Department of Medical Oncology, Dana-Farber Cancer Institute, Boston, MA, USA; Broad Institute of Harvard and MIT, Cambridge, MA, USA; Harvard Graduate Program in Biological and Biomedical Sciences, Boston, MA, USA; Harvard Graduate Program in Bioinformatics and Integrative Genomics, Boston, MA, USA; Department of Thoracic and Head and Neck Oncology, MD Anderson Cancer Center, Department of Genomic Medicine, MD Anderson Cancer Center; Division of Hematology/Oncology, Department of Medicine, University of California, San Francisco, San Francisco, CA 94143, USA; Parker Institute for Cancer Immunotherapy, University of California, San Francisco, San Francisco, CA 94143, USA

**Author notes:** corresponding author. Eliezer M. Van Allen, Dana-Farber Cancer Institute, 450 Brookline Ave, Boston MA 02215.

## Abstract

Given the growing number of clinically integrated cancer single-cell transcriptomic studies, robust differential enrichment methods for gene signatures to dissect tumor cellular states for discovery and translation are critical. Current analysis strategies neither adequately represent the hierarchical structure of clinical single-cell transcriptomic datasets nor account for the variability in the number of recovered cells per sample, leading to results potentially confounded by sample-driven biology with high false positives instead of accurately representing true differential enrichment of group-level biology (e.g., treatment responders vs. non-responders). This problem is especially prominent for single-cell analyses of the tumor compartment, because high intra-patient similarity (as opposed to inter-patient similarity) results in stricter hierarchical structured data that confounds enrichment analysis. Furthermore, to identify signatures which are truly representative of the entire group, there is a need to quantify the robustness of otherwise statistically significant signatures to sample exclusion. Here, we present a new nonparametric statistical method, BEANIE, to account for these issues, and demonstrate its utility in two cancer cohorts stratified by clinical groups to reduce biological hypotheses and guide translational investigations. Using BEANIE, we show how the consideration of sample-specific versus group biology greatly decreases the false positive rate and guides identification of robust signatures that can also be corroborated across different cell type compartments.

## Introduction

Single-cell transcriptomic profiling of patient tumors has enabled high-resolution dissections of disease progression and treatment response. Building on seminal cellular atlases for specific cancer types, many studies are increasingly focused on deriving hypotheses by evaluating groups of patients (e.g., treated vs. untreated, responders vs. non-responders, and early-vs. late-stage) for differences in gene signatures (which may be an experimentally and/or computationally derived aggregation of related genes or pathways) between the two groups. For this purpose, methods such as the Mann-Whitney U (MWU) tests and Generalised Linear Models (GLMs) have been conventionally used in bulk RNA-sequencing (bulk RNA-seq) studies as well as single-cell transcriptomic analyses;^1–4^ however, they may have a number of limitations for the latter application. First, these methods assume mutual independence of samples, and although this is not problematic for bulk RNA-seq analyses, cells derived from the same patient in single-cell analyses do not satisfy this criteria. Second, these methods fall short of representing the hierarchical structure of tumor single-cell transcriptomic data, as tumor cells tend to exhibit more intra-patient similarity as compared to inter-patient similarity due to the expression of patient-specific transcriptional programs driven by DNA-level alterations and epigenetics.^5–8^ This challenge, in turn, may lead to differential enrichment results being skewed by patient-specific biology, instead of representing genuine group biology. Finally, the number of cells (and hence, data points) sequenced in these single-cell transcriptomic datasets are typically large compared to bulk RNA-seq datasets, thereby potentially increasing the power of statistical tests to detect differences (by rejecting the null hypothesis) between the groups under consideration, which may not reflect biologically or clinically relevant observations. These challenges exist for other cell types as well, including the immune and stromal cells, albeit to a lesser extent. As a result of these significant methodological challenges, single-cell transcriptomic case/control analyses of cancer samples have thus far often not involved detailed assessments of the tumor compartments, which has restricted the capability to learn from tumor cellular programs in increasingly complex clinical contexts.

To maximize the utility of single-cell transcriptomic analyses between clinically relevant patient populations and determine how tumor cell programs differ between groups of patients, we developed a nonparametric statistical group biology estimation method (group **B**iology **E**stim**A**tion i**N** s**I**ngle c**E**ll, “BEANIE”) inspired from *He et al.*,^9^ addressing the above-mentioned issues (Fig. 1, see Methods). This method first estimates the statistical significance (empirical p-value) of the test signatures through a Monte Carlo approximation of the test signatures’ p-value distribution (test distribution) and that of the random signatures’ p-value distribution (background distribution), followed by contextualisation of the former with respect to the latter. It then uses the leave-one-out cross-validation approach (sample exclusion) to infer robustness of the gene signatures (see Methods). We used publicly available datasets to demonstrate the utility of this method, and present suggested guidelines for the design of clinically embedded single-cell transcriptomic studies in oncology.

**Figure 1.**
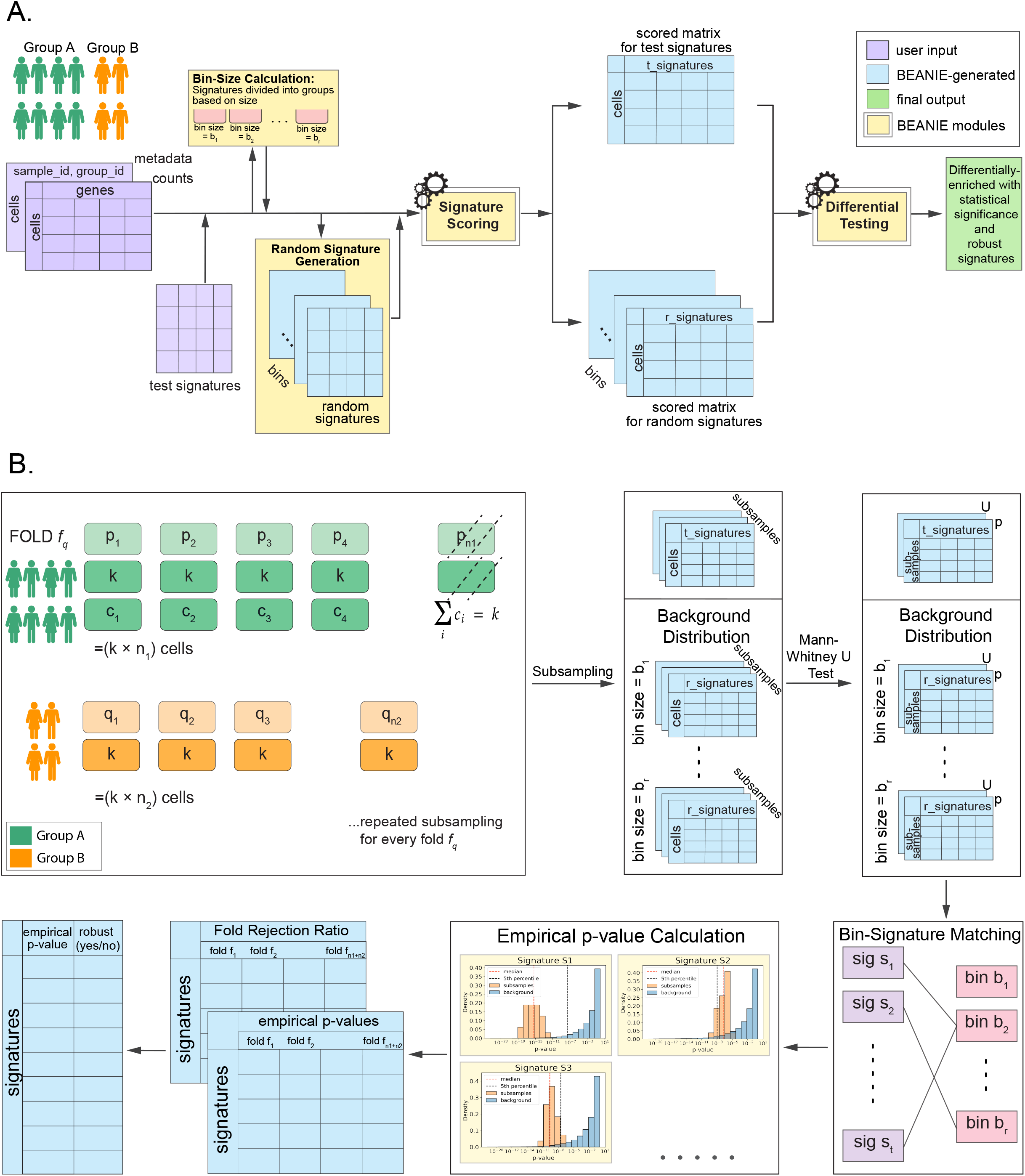
Overview of the BEANIE method. A. Overall workflow: A counts matrix, sample IDs, group IDs, and a list of signatures for which differential enrichment will be tested (test signatures, t_signatures) are provided as user input. Based on the gene set size, test signatures are first divided into bins. For each bin size, a list of random signatures (r_signatures) of the same gene set size is generated, to be later used for p-value calculation and biological interpretation. Signature scoring per cell is performed for both random signatures and test signatures, followed by differential enrichment testing. B. Differential enrichment testing workflow: The differential enrichment testing algorithm is based on a combination of Monte Carlo approximation of empirical p-value through subsampling, and leave-one-out cross validation through sample exclusion. The data is first divided into folds, where each fold f_q_ represents the exclusion of a sample from either of the comparison groups. This is followed by the subsampling step, where an equal number of cells are subsampled from every sample to ensure equal patient representation. Next, a Mann-Whitney U test is performed per subsample for all folds, for both the test signatures and the background distribution (generated from the random signatures). The test signatures are then matched to their corresponding background distribution based on bin size, and an empirical p-value (percentile of the test distribution’s median with respect to the background distribution) is calculated per test signature for every fold f_q_. Additionally, a Fold Rejection Ratio (FRR) (see Methods) is calculated per test signature for every fold, and is used to determine the overall robustness of the test signature to sample exclusion.

## Results

We evaluated single-cell transcriptomic data in two cancer types (melanoma and lung cancer) that have the following clinical groups for comparison: (i) response to treatment; and (ii) disease progression.^1, 2, 10–12^ We contextualised their tumor compartments with signatures from the Molecular Signatures Database (MSigDB),^13, 14^ including Hallmark (n = 50) and Oncogenic (n = 189) gene sets. We compared results obtained from MWU tests followed by Benjamini-Hochberg (BH) corrections and GLMs with results obtained from BEANIE, and characterised our approach relative to these methods (Table 1). Details regarding the implementation and comparisons are available in the Methods section.

**Table 1.** Number of differentially enriched signatures identified with the three methods (MWU test, GLMs, and BEANIE) using Hallmark (n = 50) and Oncogenic (n = 189) gene sets.

|  |  | MWU test | GLM | BEANIE |
| --- | --- | --- | --- | --- |
| **Melanoma (Immune Checkpoint Blockade (ICB)-naive vs. ICB-exposed)** | Hallmark gene sets | 45/50 | 27/50 | 5/50 |
|  | Oncogenic gene sets | 166/189 | 58/189 | 9/189 |
| **Lung (Early-stage vs. Late-stage)** | Hallmark gene sets | 46/50 | 44/50 | 3/50 |
|  | Oncogenic gene sets | 168/189 | 128/189 | 3/189 |
| **Lung (Tyrosine Kinase Inhibitor (TKI) - naive vs. TKI-exposed)** | Hallmark gene sets | 47/50 | 33/50 | 2/50 |
|  | Oncogenic gene sets | 152/189 | 82/189 | 1/189 |

### Group biology analysis of Immune Checkpoint Blockade-naive vs. -exposed melanoma

We first evaluated a melanoma dataset,^1^ which included data for both immune checkpoint blockade (ICB)-naive and ICB-exposed patients, to assess methodologies for comparing clinical treatment states in tumor cells. All of the ICB-exposed samples were resistant to treatment and were biopsied from the metastatic sites. We excluded samples having less than 50 tumor cells, and, in total, there were 1891 tumor cells across 14 patients, with 7 patients per group (Fig. S1, see Methods). We first assessed the data with Hallmark and Oncogenic gene set signatures from MSigDb, to characterise treatment-driven biology within the tumor compartment (Fig. 2, Table 1).

**Figure 2.**
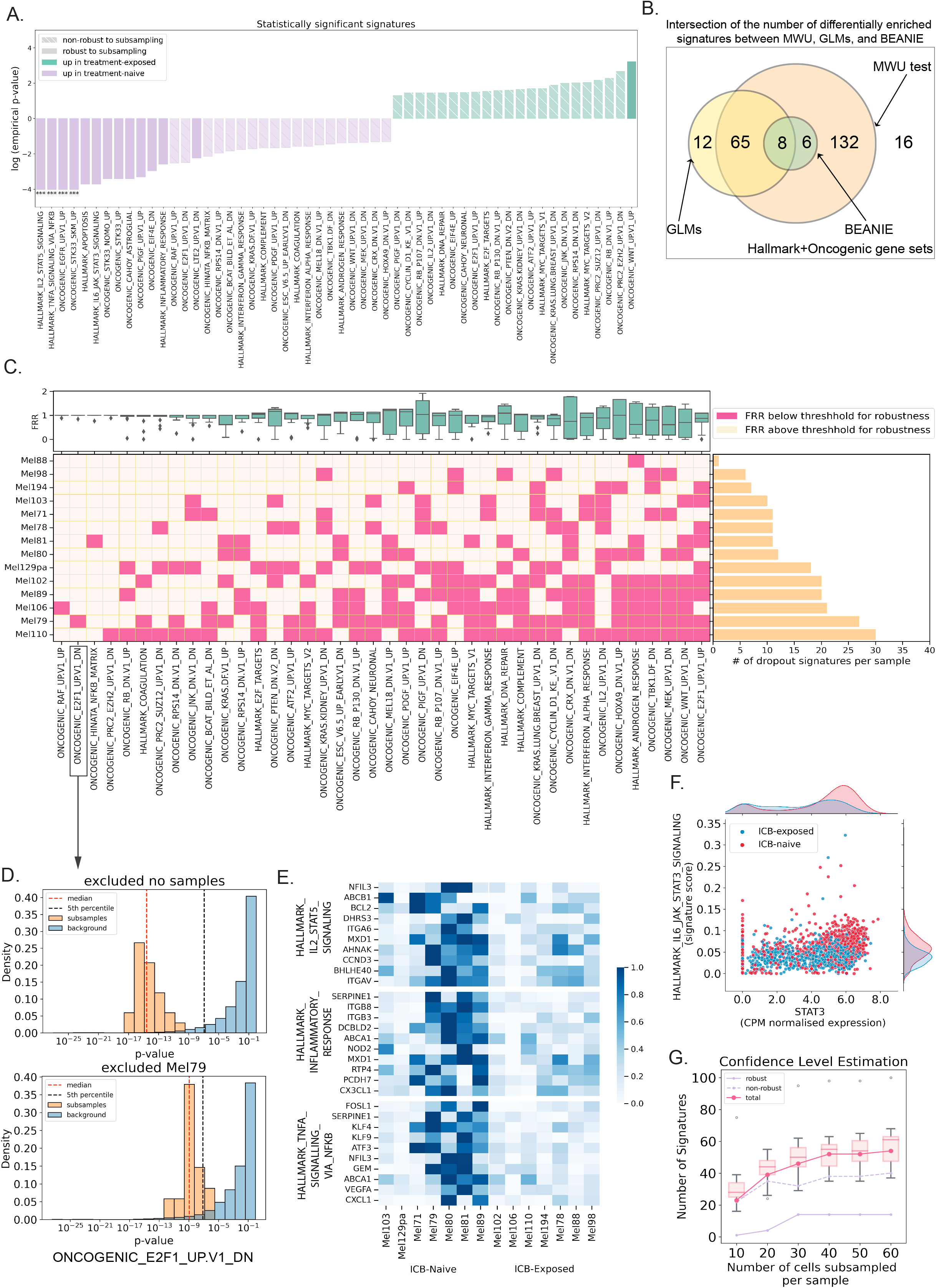
Group biology analysis of the tumor compartment from ICB-naive vs. ICB-exposed melanoma patient samples. A. Bar plot displaying the log(empirical p-value) for all of the signatures identified as statistically significant (empirical p-value ≤ 0.05), along with their robustness status. A ‘***’ above a bar indicates that the empirical p-value for that test signature was below the minimum empirical p-value measured. B. Venn diagram quantifying the intersection of signatures identified as differentially enriched with statistical significance by the three methods (MWU test with a BH correction, GLMs, and BEANIE). C. Plot depicting the signatures identified as statistically significant but non-robust to sample exclusion by BEANIE, the distribution of their Fold Rejection Ratios (FRRs), and the sample IDs having FRRs less than the threshold used to determine robustness, along with a horizontal bar plot of the number of statistically significant but non-robust signatures (dropout signatures) per sample. D. Histogram illustrating the sample exclusion procedure implemented within BEANIE shifting the test distribution to the right such that it overlaps with the background distribution, leading to the fold’s empirical p-value being greater than 0.05. E. Heatmap revealing the differential top constituent genes (ranked according to log2 fold change and robustness) from three of the statistically significant and robust signatures identified by BEANIE across all patients from the ICB-naive group. F. Joint scatter and density plot demonstrating a positive correlation between the HALLMARK_IL6_JAK_STAT3_SIGNALING signature score and STAT3 gene expression in individual cells. G. Plot illustrating BEANIE’s stability to subsample size for the test signatures used. The curve plateaued as the number of statistically significant test signatures, irrespective of robustness status, reached saturation as the subsample size approached 60 (the max subsample size possible within the constraints of this dataset [see Methods]), whereas the curve plateaued around the subsample size of 30 as the number of signatures identified as both statistically significant and robust reached saturation.

We observed that the MWU test followed by a BH correction and GLMs predicted a large number of differentially enriched signatures (p-value ≤ 0.05), whereas BEANIE was more conservative, detecting fewer signatures as differentially enriched (Fig. 2b, Table S1). Notably, a majority of the signatures identified as significant by MWU test and GLMs were labelled as non-significant and non-robust to sample exclusion by BEANIE.

Among signatures that were identified as statistically significant and robust to sample exclusion by BEANIE (see Methods), signatures upregulated in the ICB-naive group include those for genes upregulated by *STAT5* in response to *IL2* stimulation (HALLMARK_IL2_STAT5_SIGNALING), genes regulated by NF-κB in response to TNF (HALLMARK_TNFA_SIGNALING_VIA_NFKB), and genes defining inflammatory response (HALLMARK_INFLAMMATORY_RESPONSE). We also verified a previously identified T cell exclusion signature^1^ upregulated in the ICB-exposed group as statistically significant and robust to sample exclusion with the BEANIE method. To verify gene-level differential expression, we used a MWU test and observed differential *IL2* gene expression for the ICB-naive group in the T cell compartment (p-value = 0.0078), corroborating our finding in the tumor compartment (differential HALLMARK_IL2_STAT5_SIGNALING). We also identified the top constituent genes (ranked according to log2 fold change and robustness to sample exclusion, see Methods) for these three signatures, and found that these genes were differential in the tumor compartment uniformly across samples of a given group (Fig. 2e, Table S2). Together, these results describe the tumor microenvironment of the ICB-exposed group (consisting of treatment-resistant patients) as one depleted of T cells, with reduced IL2-STAT5 signaling, TNFA-NFKB signaling, and inflammatory response relative to the ICB-naive group.

Additionally, we found that the signature for genes upregulated by *IL6* via STAT3 (HALLMARK_IL6_JAK_STAT3_SIGNALING) was upregulated in the ICB-naive group and statistically significant and robust to sample exclusion. Using a MWU test, we found differential *STAT3* expression in the tumor compartment for the ICB-naive group (p-value = 3.28e-36). Furthermore, we also found a positive correlation between the *STAT3* expression and HALLMARK_IL6_JAK_STAT3_SIGNALING signature score in the tumor cells on an individual cell basis (Fig. 2f). This observation supports the finding that *IL6* could potentially induce downstream signaling via *STAT3* in the tumor cells of the ICB-naive group.^15, 16^

We further examined the cause for non-robustness of the signatures that were identified as statistically significant but not robust to sample exclusion by BEANIE (Fig. 2c). We found that the exclusion of one or more samples led to statistically non-significant results, in contrast to when the sample was included, by shifting the empirical p-value to greater than 0.05 as a result of an overlap between the test distribution and the background distribution as shown in Fig. 2d (see Methods). For example, the signature ONCOGENIC_RAF_UP.V1_UP was not robust to the exclusion of sample Mel106, and this particular sample was also the cause of non-robustness for 21 other signatures. This variability due to sample exclusion was also not explained by any of the other available clinical variables (e.g., age, sex). Therefore, these signatures were driven by sample-specific biology, and were consequently not representative of the group-level biology, but would have otherwise been considered differentially enriched with statistical significance using either of the conventional MWU test or GLM approaches.

We next investigated the methodological stability with respect to subsample size (Fig. 2g), and accordingly repeated BEANIE’s workflow using smaller subsample sizes. We found that a smaller subsampling of cells led to fewer signatures that were identified as statistically significant and non-robust to sample exclusion, and even fewer that were identified as both statistically significant and robust to sample exclusion. However, the number of statistically significant and robust signatures identified by BEANIE reached saturation around the subsample size of 30 cells per sample, indicating that the subsample size of 60, which had been used for all of the aforementioned results, could successfully capture all statistically significant signatures from the test signature sets that were also robust to sample exclusion.

To assess the ability to detect noise from a true signal, we additionally used a curated set of immune cell surface marker signatures^17^ (including signatures for T cells, NKT cells, NK cells, B cells, mast cells, and a joint dendritic cell/macrophage signature), that should not be relevant to tumor cells, to test the performance of the three methods (MWU test with a BH correction, GLMs, and BEANIE). We observed that a MWU test with a BH correction led to a p-value ≤ 0.05 for all signatures except the B cell signature and GLMs led to a p-value ≤ 0.05 for NKT cell, B cell, NK cell, and the joint dendritic cell/macrophage signatures. By contrast, BEANIE correctly predicted all of the immune cell surface marker signatures as both statistically non-significant and non-robust to sample exclusion (Table S3).

Finally, we evaluated BEANIE’s performance on the previously reported 18 T cell exclusion signatures^1^ for the tumor cell compartments of a reduced set of patients from the original set used to derive these (only patient samples with greater than 50 tumor cells were retained, as described above). We observed that while a MWU test with a BH correction had p-values ≤ 0.05 for 18/18 signatures and GLMs had p-values ≤ 0.05 for 11/18 signatures, BEANIE had an empirical p-value ≤ 0.05 for 17/18 signatures and additionally found 10/18 of them to be robust to sample exclusion.

### Group biology analysis of distinct clinical states in non-small cell lung carcinoma

In an effort to demonstrate the applicability of BEANIE for a meta-analysis composed of multiple single-cell transcriptomic datasets, we next analyzed the tumor compartments from four published lung cancer studies^2, 10–12^ to evaluate potential differentially enriched signatures between early-vs. late-stage samples. We selected patient samples which satisfied the following criteria: (i) had more than 50 tumor cells; (ii) were classified as adenocarcinoma; (iii) were staged as either I, II, or IV (early-stage = I and II; late-stage = IV); and (iv) had received no prior treatment at the time of sample collection. Filtering according to these criteria yielded a total of 18251 malignant cells across 17 patients (11 early-stage, 6 late-stage) (Fig. 3a).

**Figure 3.**
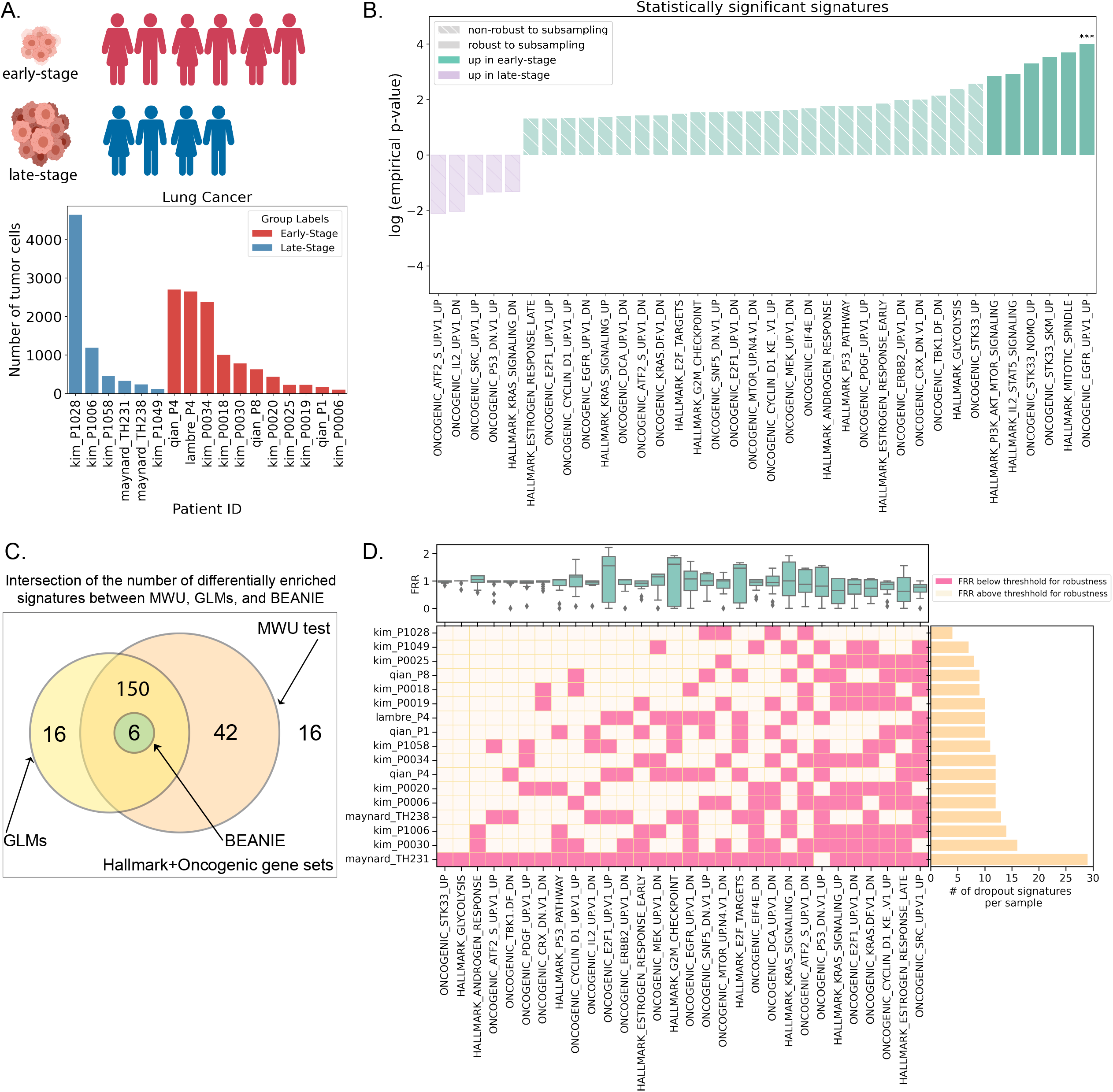
Group biology analysis for early-vs. late-stage non-small cell lung cancer. A. Overview of the integrated dataset from four studies and a bar plot depicting the number of tumor cells per patient sample. B. Bar plot displaying the log(empirical p-value) for all of the signatures identified as statistically significant (empirical p-value ≤ 0.05), along with their robustness status. C. Venn diagram quantifying the intersection of signatures identified as differentially enriched with statistical significance by the three methods (MWU test with a BH correction, GLMs, and BEANIE). D. Plot depicting the signatures identified as statistically significant but non-robust to sample exclusion by BEANIE, the distribution of their FRRs, and the patient IDs having FRRs less than the threshold used to determine robustness, along with a horizontal bar plot of the number of statistically significant but non-robust signatures (dropout signatures) per sample.

We sought to characterise the tumor compartment with Hallmark and Oncogenic gene sets from MSigDb (Fig. 3b, Table 1). Again, a large number of gene sets predicted as differentially enriched with statistical significance by a MWU test with a BH correction and GLMs were identified as statistically non-significant and non-robust to sample exclusion by BEANIE (Fig. 3c, Table S1). We found a signature composed of genes important for mitotic spindle assembly (HALLMARK_MITOTIC_SPINDLE) to be statistically significant and robust to sample exclusion for early-stage lung tumors with BEANIE, consistent with prior studies.^18^ Another signature comprised of genes encoding proteins involved in glycolysis and gluconeogenesis (HALLMARK_GLYCOLYSIS) was also found to be statistically significant and robust to sample exclusion for the early-stage tumors with BEANIE, which is in agreement with a prior study^19^ describing an association between TKI treatment and its effect on decreased activity of glycolysis. Furthermore, the top constituent genes for both of these signatures were consistently upregulated across all samples (Fig. S2, Table S2). Thus, BEANIE was able to detect both statistically significant and robust signatures in the meta-analysis of multiple single-cell transcriptomic datasets.

We next sought to evaluate the tumor compartment from two of the lung cancer datasets (Kim et al.,^10^ Maynard et al.^11^) for treatment responses to tyrosine kinase inhibitors (TKIs). We selected patient samples which satisfied the following criteria: (i) had more than 50 tumor cells; and (ii) the biopsy was derived from the primary tumor. These filtering criteria led to a total of 7576 malignant cells across 10 patients (6 TKI-naive, 4 TKI-exposed) (Fig. S3).

We again used the Hallmark and Oncogenic gene sets to characterise the tumor compartment (Fig. 4a, 4b, Fig. S3; Table 1, Table S1). Among the signatures that were found to be statistically significant and robust to sample exclusion with BEANIE, signatures upregulated in the TKI-exposed group included a signature for genes upregulated in response to *IFNG* (HALLMARK_INTERFERON_GAMMA_RESPONSE) and a signature for genes upregulated by the overexpression of *WNT1* (ONCOGENIC_WNT_UP.V1_UP). We identified the top constituent genes of both signatures, and found them to be consistently upregulated across all samples in the TKI-exposed group (Fig. 4c, Table S2). Interferon gamma response has been described to be associated with response to TKI treatment in non-small cell lung cancers.^20^ Using a MWU test, we observed that genes encoding the *IFNG* receptors (*IFNGR1, IFNGR2*) were differentially expressed with statistical significance in the tumor cells of the TKI-exposed group (p-value [*IFNGR1*] = 3.22e-104, p-value [*IFNGR2*] = 1.81e-13, Fig. S3). WNT signaling has also been extensively studied in the context of cancer development, and increased WNT signaling has been associated with tumor progression and metastasis in many different cancers.^21^ We assessed potential intratumoral differential gene expression of *WNT1* in the tumor cells and found an absence of intratumoral *WNT1* expression altogether. We then assessed potential *WNT1* differential gene expression in specific immune cell compartments (NK cells, macrophages, and T cells) and found a statistically significant differential expression of *WNT1* for the TKI-exposed group within the T cell compartment (MWU test, p-value = 8.77e-18), indicating putative cross-compartment communication between the T cells and tumor cells via *WNT1* signaling. To further validate this, we used a MWU test to investigate possible differential gene expression of *WNT1* receptors (*FZD1, FZD2*) in the tumor compartment and found both of the receptors to be upregulated in the TKI-exposed group (p-value [*FZD1*] = 6.35e-21, p-value [*FZD2*] = 1.28e-27). Of note, patients who were treated with TKI were classified with RECIST as having either PD (Progressive Disease) or RD (Residual Disease), which raises the hypothesis that these patients may have developed therapeutic resistance through the WNT/beta-catenin signaling pathway in alignment with prior preclinical studies.^22^

**Figure 4.**
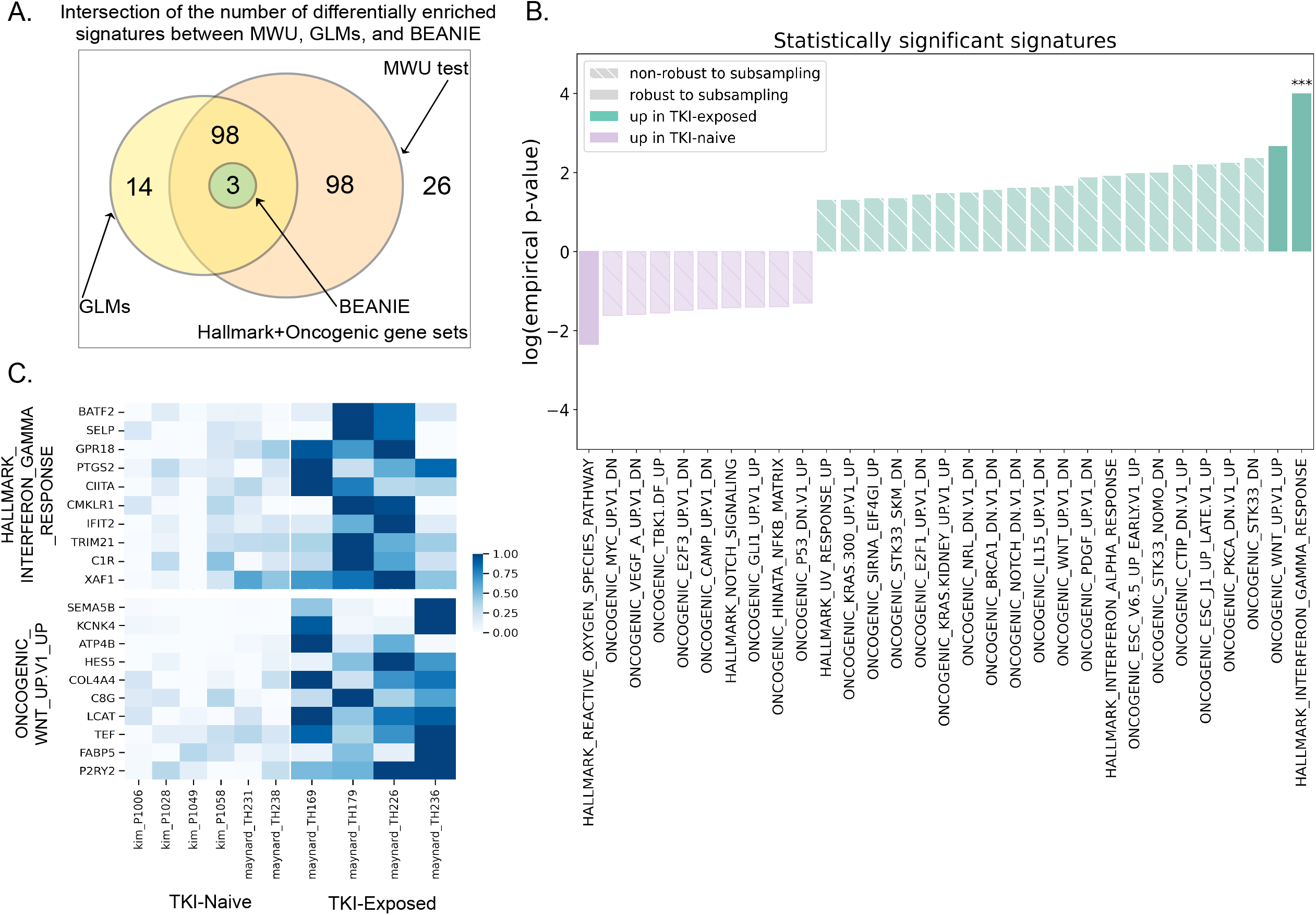
Group biology analysis for TKI-naive vs. TKI-exposed non-small cell lung cancer. A. Venn diagram quantifying the intersection of signatures identified as differentially enriched with statistical significance by the three methods (MWU test with a BH correction, GLMs, and BEANIE). B. Bar plot displaying the log(empirical p-value) for all of the signatures identified as statistically significant (empirical p-value ≤ 0.05) along with their robustness status. A ‘***’ above a bar indicates that the empirical p-value for that test signature was below the minimum empirical p-value measured. C. Heatmap showing the differential top constituent genes (ranked according to log2 fold change and robustness) from the HALLMARK_INTERFERON_GAMMA_RESPONSE and ONCOGENIC_WNT_UP.V1_UP signatures across all samples from the TKI-exposed group.

In addition, we estimated the stability of BEANIE to subsample size for the test signatures used (Fig. S3), and found that the number of robust signatures identified persistently increased at the maximum sample size, indicating the possibility that some of the signatures classified as robust could have been instead classified as non-robust. This may be a result of unbalanced samples per group being tested or may demonstrate the necessity of additional biological samples for the clinical context being evaluated.

### Estimation of the False Positive Rate

In order to estimate the chance of occurrence of incorrectly identified statistically significant and robust signatures with BEANIE, we calculated the false positive rate (type I error) (see Methods, Fig. 5) for all three methods (MWU test with a BH correction, GLMs, BEANIE) and clinical contexts (ICB-naive vs. -exposed melanoma, early-vs. late-stage lung cancer, and TKI-naive vs. -exposed lung cancer) for the signatures that had been classified by BEANIE as both statistically significant and robust to sample exclusion.

**Figure 5.**
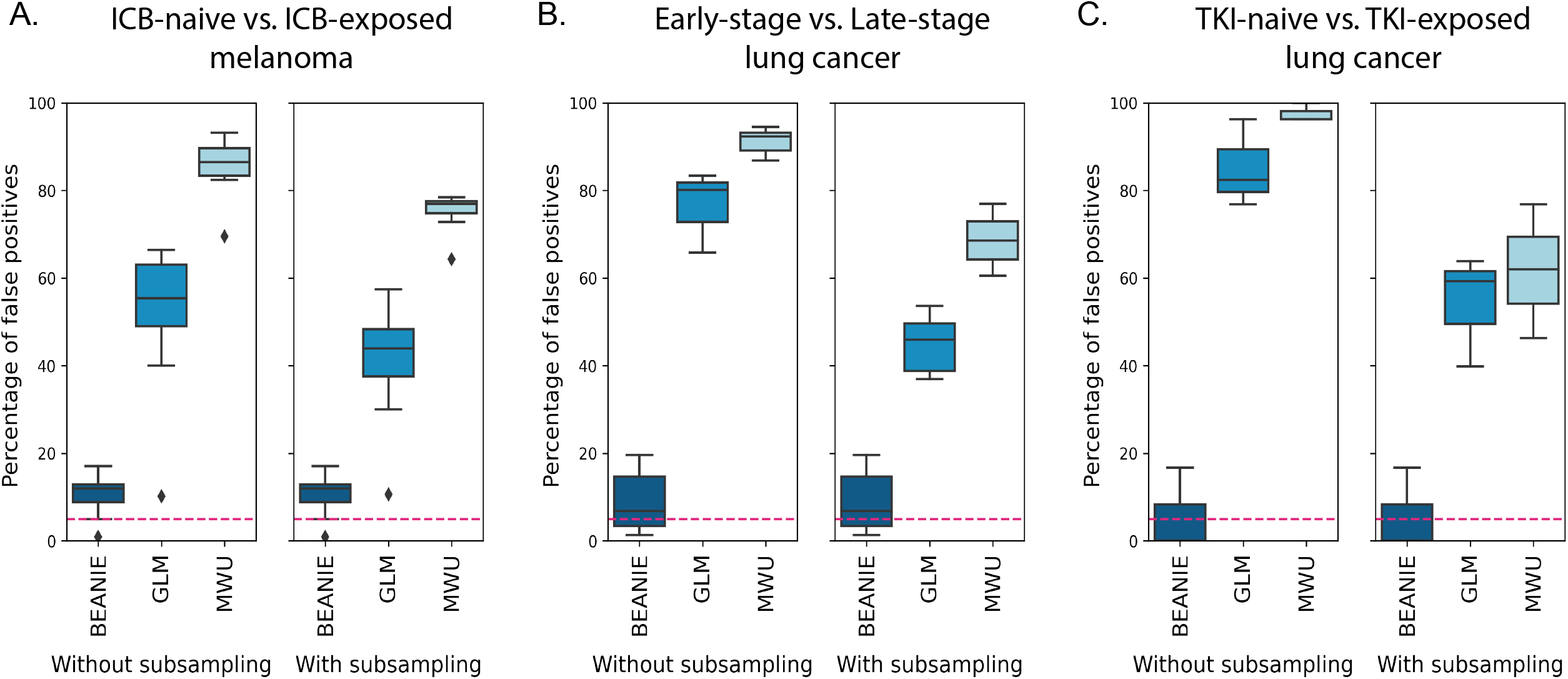
False positive rate (type I error) for the three methods: BEANIE, GLMs, and a MWU test followed by a BH correction. The false positive rate for signatures (from Hallmark and Oncogenic gene sets) which were classified as statistically significant and robust to sample exclusion by BEANIE for: A. ICB-naive vs. ICB-exposed melanoma dataset, B. Early-stage vs. Late-stage lung cancer dataset, and C. TKI-naive vs. TKI-exposed lung cancer dataset The dashed pink line denotes the 5% error mark.

We observed that across all datasets, the MWU test with a BH correction had a high average false positive rate, followed by GLMs which exhibited a moderately high average false positive rate. By contrast, BEANIE had the lowest average false positive rate, that in some cases also approached the significance level (alpha) of 5% (Table 2). Individual false positive rates calculated for all robust and statistically significant signatures can be found in Table S4.

**Table 2.** Average false positive rate for the three datasets (ICB-naive vs. ICB-exposed melanoma, early-stage vs. late-stage lung cancer, and TKI-naive vs. TKI-exposed lung cancer) across the three methods (MWU test with a BH correction, GLMs, and BEANIE).

|  | MWU test + BH correction |  | GLMs |  | BEANIE |
| --- | --- | --- | --- | --- | --- |
|  | Without subsampling | With subsampling | Without subsampling | With subsampling |  |
| **ICB-naive vs. ICB-exposed** | 85.74% | 75.57% | 52.58% | 41.72% | 10.63% |
| **Early-stage vs. Late-stage** | 91.23% | 68.6% | 77.03% | 44.93% | 9.01% |
| **TKI-naive vs. TKI-exposed** | 97.53% | 61.72% | 85.18% | 54.32% | 5.55% |

To evaluate how a smaller number of cells being tested, and thereby reduced statistical power, would impact the false positive rate for a MWU test with a BH correction and GLMs, we subsampled cells from each sample being tested to a number equivalent to BEANIE’s subsample size and repeated the type I error estimation. We found that subsampling decreased the false positive rates for both a MWU test with a BH correction and GLMs, but that their false positive rates were still relatively higher than those calculated with BEANIE, corroborating BEANIE’s aptitude for detecting robust and true signals as compared to the other two methods, and also reinforcing the need to incorporate robustness estimation into differential enrichment testing.

### Group biology estimation in immune cells

While this strategy was primarily developed to overcome challenges for differential enrichment testing specifically within the tumor cell compartment, we also evaluated whether the subsampling and sample exclusion approach implemented within BEANIE would likewise yield biological insights in immune cell compartments, as well as to further validate some of the initial hypotheses from the tumor compartment analyses. As a test case, in continuation of the preliminary evaluation of the CD8+ T cell compartment as described in the earlier tumor compartment analysis, we more comprehensively dissected the CD8+ T cell compartment in ICB-naive vs. -exposed melanoma patients in an isolated context here (Fig. S1).

We filtered out samples which had fewer than 50 CD8+ T cells, yielding a total of 1292 cells across 11 patients (5 ICB-naive, 6 ICB-exposed). We evaluated the potential statistically significant differential enrichment and robustness to sample exclusion of various signatures representing a range of CD8+ T cell subtypes and states from Oliveira et al.^23^ between the ICB-naive and - exposed patient groups. We found that previously reported signatures for early activated CD8+ T cells (Sade-Feldman_5^24^) and memory precursor effector CD8+ T cells (Joshi_MPEC,^25^ murine-derived) were statistically significant and robust to sample exclusion, and upregulated for the ICB-naive patient group. This result substantiates the finding from the tumor compartment of the same dataset, where we had identified higher IL2-STAT5 signaling, TNF activation, and inflammatory response in the tumor cells from the ICB-naive group as described, which may be a result of CD8+ T cell activation in the naive condition.

Therefore, in addition to the demonstrated utility for tumor cell compartments, BEANIE likewise exhibited capacity for cross-compartment validations of group biology in single-cell non-tumor populations as well.

## Conclusions

Conventional differential enrichment methods, such as a MWU test with a BH correction and GLMs, are limited in correctly estimating differential biology in clinical tumor single-cell transcriptomic datasets in two aspects. First, they have an appreciably high false positive rate, which can be attributed in part to an increased power of statistical tests (due to high cell numbers) to detect differences between groups. However, increased power does not necessarily signify a biologically relevant difference. Consequently, interpretation of these differences in a group biology context is requisite to correctly distinguish genuine group biological differences from technical artifacts (such as variation in cell numbers). We also observed that subsampling alone is insufficient to tackle this problem, and it is important to use a background distribution for contextualisation. Second, conventional differential enrichment methods do not assess the robustness of a signature to sample exclusion, and as a consequence, these methods may lead to results being sample-driven and of uncertain translational importance. This issue is particularly relevant in clinical contexts, and especially for tumor cell compartments which demonstrate higher intra-patient similarity than inter-patient similarity, as hypotheses based on group comparison (about treatment effects, disease progression, etc.) may impact future clinical trials.

To address the shortcomings of conventional differential enrichment methods, we developed BEANIE, a nonparametric statistical method for estimating group biology in clinical single-cell transcriptomic datasets. We demonstrated its application on publicly available datasets from six clinical single-cell transcriptomic studies, and illustrate its aptitude to detect statistically significant and robust gene signatures as compared to conventional methods, through its low false positive rate as compared to its counterparts (MWU test followed by a BH correction and GLMs). We illustrated its extensive application in the tumor compartment, and its potential utility for the immune compartment as well. It may likewise be used to identify differential enrichment of gene signatures in the stromal compartment. Finally, we demonstrated that BEANIE is adept at distinguishing sample-driven signatures from group-driven signatures, whereas conventional differential enrichment methods fail to do so. Alternate models for representation of tumor single-cell data include hierarchical linear models; however, unlike BEANIE, they are parametric and therefore assume normality and homogeneity of variance for the data.

Despite its potential to estimate group biology and pinpoint both statistically significant as well as robust and therefore prospective biologically relevant signatures in single-cell transcriptomic dissections, BEANIE also possesses a few limitations. First, in spite of its demonstrated value in single-cell transcriptomic tumor compartment analyses, BEANIE’s widespread applicability in the immune compartment may be limited, in part due to an absence of comprehensive databases with precise and rigorous signatures representing discrete cell types, states, and pathways. In fact, the ultimate utility of BEANIE’s or any group biology analysis tool’s framework is in part contingent on the quality of the gene signatures being tested, including for the tumor compartment. Moreover, there also exists scope to further improve the false positive rate within the BEANIE method. In addition, we do not currently have an understanding of why some patient samples are more prone to contribute to the non-robustness of certain signatures as compared to other patient samples, and having additional clinical information (e.g., mutational status) could potentially help delineate some of the biology behind this. Lastly, despite the ability to estimate group biology and identify statistically significant and robust signatures between patient groups with BEANIE, current clinical single-cell transcriptomic datasets have an overall small sample size, which indicates that they are likely not an adequate representation of the broader population and hence could lead to introduction of false negatives (type II error). Therefore, in general, larger datasets, such as those generated via consortium efforts, are needed to improve our ability to draw robust conclusions, and minimise putative false negatives. Broadly, dedicated efforts to analyze larger clinically integrated single-cell cohorts that reflect the diverse clinical and therapeutic contexts across cancer types will accelerate our understanding of the cell states that promote treatment resistance for translational discovery.

## Methods

### Data Preprocessing

#### Melanoma Dataset

We selected cells which were labelled as malignant (authors made use of inferCNV^28^ to identify malignant cells).

#### Lung Cancer Datasets

Owing to the variability in collected datasets from the four studies (Kim et al.^10^, Maynard et al.^11^, Qian et al.^12^, and Lambrechts et al.^2^), we carefully assessed the metadata files available. For Lambrechts et al., we reached out to the authors to acquire their Seurat object containing patient ID and cell ID labelling. We used the following criteria for the selection of cells for analysis: (i) must be of epithelial origin; (ii) must be identified as malignant by the authors (all studies made use of inferCNV^28^ to identify malignant cells); and (iii) must be isolated from the primary site (i.e., lung). We also removed cells from patients that had locally advanced lung cancer (stage III tumors), as they are more difficult to classify into early-versus late-stage.^26^

### BEANIE’s Workflow

#### Preprocessing and Normalisation

The raw counts matrix is normalised by the library size and converted to counts per million (CPM normalization) to account for differences in library sizes of different cells. Pre-normalised matrices may also be used, in which case this step is ignored. Genes with no expression across all cells are excluded.

#### Signature Scoring

For each cell *c_i_* in the normalised counts matrix, signature scoring is performed for the set of gene signatures provided as input by the user (test signatures). The default signature scoring method is adapted from AUCell.^27^

i. For each gene *g_k_*, the cells are ranked by calculating the percentile of each cell across the gene *g_k_* in terms of normalised expression of the gene, i.e., cells with higher expression values of that particular gene will have a higher percentile. The ties are randomly broken (i.e. if two cells have the exact same expression of the gene, which is common in single-cell datasets, those cells are randomly assigned a percentile value).

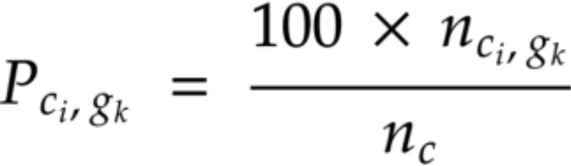

where 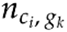 = ordinal rank of *c_i_* for expression of *g_k_* (sorted from smallest to largest), 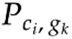 = percentile of *c_i_* for expression of *g_k_*, and *n_c_* is the total number of cells
ii. Next, for every cell *c_i_*, genes are ranked based on their calculated percentile values across that cell. Genes which have a higher percentile across the cell are given lower ranks. This scoring system takes into account the importance of each gene in a given cell relative to that gene’s importance in other cells, i.e., genes which have a lower rank are more important for the cell in question as compared to genes with a higher rank.

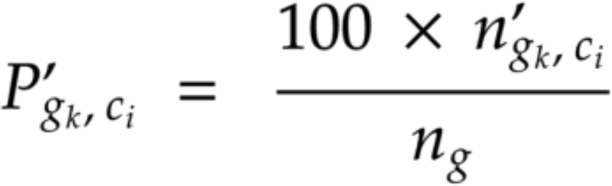

where 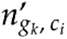 = ordinal rank of *g_k_* for 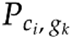 values (sorted from largest to smallest), 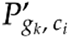 = percentile of *g_k_* for 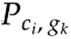 values, and *n_g_* is the total number of genes for the cell *c_i_*
iii. For each gene signature *S_j_*, a recovery curve per cell *c_i_* is generated by calculating the enrichment of the top constituent genes ranked from *S_j_,* followed by a calculation of the Area

Under the Curve (AUC), which measures the expression of *c_i_’*s top constituent genes ranked from *S_j_*. The AUC is therefore the score of the cell for *S_j_*.

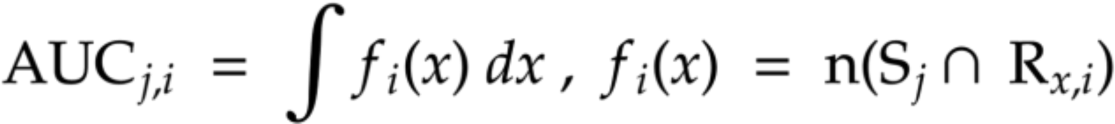

where *S_j_* = set of genes comprising a gene signature

and *R_x,i_* = set of top constituent *x* genes based on 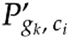

Other signature scoring methods available in BEANIE include weighted mean and z-scoring.

## Background Distribution Generation

A background distribution is generated for the biological interpretability of the results as follows: (i) Bins are created based on the gene set size of each signature *S_j_* (default bin size = 10, tunable parameter). (ii) Random signatures (r_signatures) (R_k_, k = 1, 2, …, n_b_, where n_b_ = total number of bins) for each of the bin sizes are generated such that they are representative of both lowly expressed and highly expressed genes. For this step, the normalised matrix is used and the genes are sorted based on their expression values across all samples. Equal numbers of genes from every 20^th^ percentile are then randomly subsampled such that the sum of all genes equals the bin size. This random sampling is repeated multiple times to generate different random signatures (*R_kl_,* l = 1, 2, …, n_r_, where n_r_ = the total number of times subsampling is repeated). The rationale for generating the random signatures is that they should not represent any biologically meaningful gene signature, and as a consequence, their differential expression can be used as a null distribution (background distribution) for interpretation of the results in a biological context. (iii) Each cell *c_i_* is scored for *R_kl_*’s using the aforementioned signature scoring method.

### Folds and Subsampling

To accomplish BEANIE’s two-fold aim of having equal sample representation and quantifying robustness for *S_j_*s, two statistical techniques, Monte Carlo approximations (subsampling) and leave-one-out cross-validation (sample exclusion), are coupled. First, the data is divided into folds (*f_q_*, q = 1, 2, …, *n_p_*, where *n_p_* = number of samples), with each fold *f_q_* representing the exclusion of one sample from either group. For each fold *f_q_*, cells are subsampled such that each sample is represented by an approximately equal number of cells. This is done by first subsampling an equal number of cells from all samples, followed by additional subsampling in the sample-excluded group to compensate for the cells that would have otherwise been subsampled from the excluded sample. The additional subsampling ensures that the total number of cells subsampled from the two groups being tested always remains constant regardless of which group the excluded sample belongs to, which is necessary to ensure that the folds are comparable with each other. The subsampling is then repeated multiple times to establish adequate representation of each patient sample.

### Identification of Differentially Enriched Signatures

A multi-step strategy is adopted to identify differentially enriched signatures. First, for each subsample belonging to the fold *f_q_*, a MWU test is performed between the two groups for every *S_j_*. Additionally, for each fold *f_q_*, a null p-value distribution is generated by a MWU test between the two groups for every *R_kl_*. The null distribution generated is fold-specific to ensure that the sample excluded from the fold is also excluded for the generation of the null distribution. The percentile of the subsample’s p-value against the null p-value distribution is then calculated, hereafter referred to as the empirical p-value. A median empirical p-value is calculated for these subsamples to represent the p-value for a given fold, followed by a median across all folds to represent the cell’s p-value. To quantify the robustness of *S_j_* to sample exclusion, a ratio (henceforth referred to as the Fold Rejection Ratio (FRR)) is defined, and calculated for every fold *f_q_*.

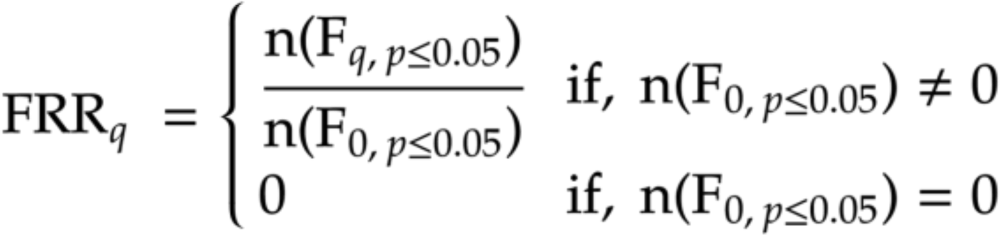

where F*_q_* = set of subsamples for the fold *f_q_* which have an empirical p-value ≤ 0.05 and F*_0_* = set of subsamples for the fold *f_0_* (when no sample is excluded) which have an empirical p-value ≤ 0.05 A FRR value closer to 1 indicates that exclusion of the sample has no effect on the empirical significance of the gene signature *S_j_*, and a lower value indicates the opposite. We use a threshold of 0.9 (hyperparameter) to call signatures as robust or not, i.e., if the FRR for a particular *S_j_* is greater than 0.9 for *all* folds, then the gene signature is considered robust to sample exclusion.

### Gene Ranking

For every gene signature *S_j_*, the genes are then ranked for the robustness of their log2 fold change between the two groups. This is particularly useful for larger gene sets. For every sample, a mean gene expression (MGE) is calculated for every gene using the normalised counts. A similar strategy of subsampling coupled with sample exclusion is used for ranking. The MGE matrix is then divided into folds, with each fold representing the exclusion of one sample. A lo g2 fold change is then calculated for each fold, and the standard deviation, along with the mean across folds, is also calculated. Genes with both outlier MGE values and outlier log2 fold changes (i.e., MGE values and log2 fold changes more than 1.5 times the interquartile range above the third quartile or below the first quartile) are classified as non-robust to sample exclusion. The final ranking of genes is performed based on decreasing log2 fold change, increasing standard deviation, and robustness status.

### Mann-Whitney U tests

Mann-Whitney U (MWU) tests followed by Benjamini-Hochberg (BH) correction are performed for the calculation of p-values. The Python package *scipy* is used for the MWU p-value calculation and the function *multipletests* from the Python package *statsmodels* is used for the BH correction.

### Generalised Linear Models

Generalised linear models (GLMs) with a binomial distribution link function are used for calculation of p-values. The Python package *statsmodels* is used to implement this method. The signature scores are used as covariates (exog variable), and the group labels (e.g., treatment-naive or - exposed and early-stage or late-stage) as the response variable to be modelled (exog variable).

### Calculation of False Positive Rate (Type I Error)

False positive rate (type I error) refers to the probability of detecting a result by chance. To calculate this, we permute the patient ID and group label in such a way that roughly equal numbers of samples from the original group labels are placed in both comparison groups. We then repeat the BEANIE workflow on these permuted datasets for signatures which are classified by BEANIE as statistically significant and robust to sample exclusion in the original dataset to evaluate the type I error rates for our predictions. In addition, we also run a MWU test followed by a BH correction and GLMs for these signatures to compare the type I error rates across the three methods. Finally, to investigate whether Monte Carlo subsampling (with equivalent statistical power to that of BEANIE’s workflow) would affect the false positive rate. For this, we subsample a random set of cells equal to the number of cells subsampled in the BEANIE workflow and repeat the MWU test and GLM methods.

For the ICB-naive vs. ICB-exposed melanoma dataset (14 samples, 7 in each group) and early-stage vs. late-stage lung cancer dataset (17 samples, 11 early-stage and 6 late-stage), we ran 1000 simulations per gene signature with the above workflow to estimate the false positive rate. For the TKI-naive vs. TKI-exposed lung cancer dataset (10 samples, 6 TKI-naive and 4 TKI-exposed), we ran 100 simulations per gene signature (due to limited combinations of equidistributed samples per group possible).

### Data Availability

All datasets used in the study are publicly available. Hallmark and Oncogenic gene sets are available for download from MSigDb.

### Code Availability

Code is publicly available as a downloadable Python package from: https://github.com/sjohri20/beanie.

## Acknowledgements

This project was funded by NIH U01 CA233100 (E.M.V.A., L.F.), R01CA227388 (E.M.V.A.), U2CCA233195 (E.M.V.A.), PCF-Movember Challenge Award (E.M.V.A.)

## Disclosures

E.M.V.A. has received research support (to institution) from Novartis and BMS. E.M.V.A. serves as a consultant or on scientific advisory boards of Tango Therapeutics, Genome Medical, Invitae, Enara Bio, Janssen, Manifold Bio, Monte Rosa. E.M.V.A. has equity in Tango Therapeutics, Genome Medical, Syapse, Enara Bio, Manifold Bio, Microsoft, Monte Rosa. E.M.V.A. receives travel reimbursements from Roche/Genentech. E.M.V.A. has filed institutional patents on chromatin mutations and immunotherapy response, and methods for clinical interpretation, and has intermittent legal consulting on patents for Foaley & Hoag.

L.F. has received research support (to institution) from Roche/Genentech, Abbvie, Bavarian Nordic, Bristol Myers Squibb, Dendreon, Janssen, Merck, and Partner Therapeutics. L.F. has served on the scientific advisory boards of Actym, Allector, Astra Zeneca, Atreca, Bioalta, Bolt, Bristol Myer Squibb, Immunogenesis, Merck, Merck KGA, Nutcracker, RAPT, Scribe, Senti, Soteria, TeneoBio, and Roche/Genentech.

M.X.H. has been a consultant to Amplify Medicines, Ikena Oncology, and Janssen. He is also currently an employee of Genentech/Roche.

J.C. has been a consultant to Tango Therapeutics. He is also currently an employee of PathAI.

N.I.V. serves on advisory boards of Sanofi and Oncocyte.

## Author Contributions

M.X.H. and E.M.V.A. conceived the original idea. S.J. developed the idea further, designed experiments, performed the analyses, and developed the Python package for the presented method. K.B., B.M.T., J.F. and M.X.H. helped in development of the method from a single-cell perspective. M.X.H., J.C. and D.L. provided input from a statistical perspective. J.P.C. provided input for the development of visualisation modules in the package. N.I.V. and D.L. helped in clinical interpretation of the results. Z.F., J.P., L.F., D.L., and E.M.V.A. contributed to the overall analyses. S.J., B.M.T., K.B., and E.M.V.A. wrote the manuscript. All authors reviewed and approved the final manuscript.

## Competing Interests statement

The authors declare no competing interests.

## Supplementary figures

**Figure S1.**
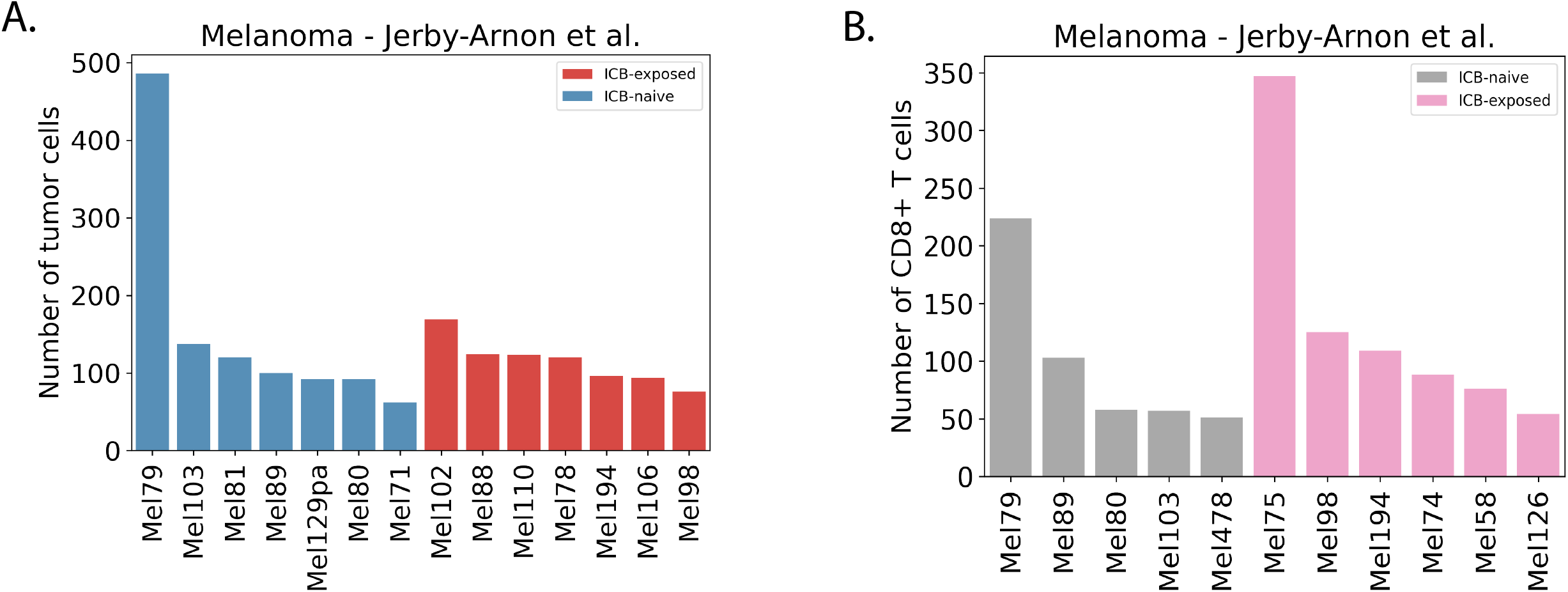
Distribution of cells from ICB-naive vs. ICB-exposed melanoma patient samples. A. Distribution of tumor cells. B. Distribution of CD8+ T cells.

**Figure S2.**
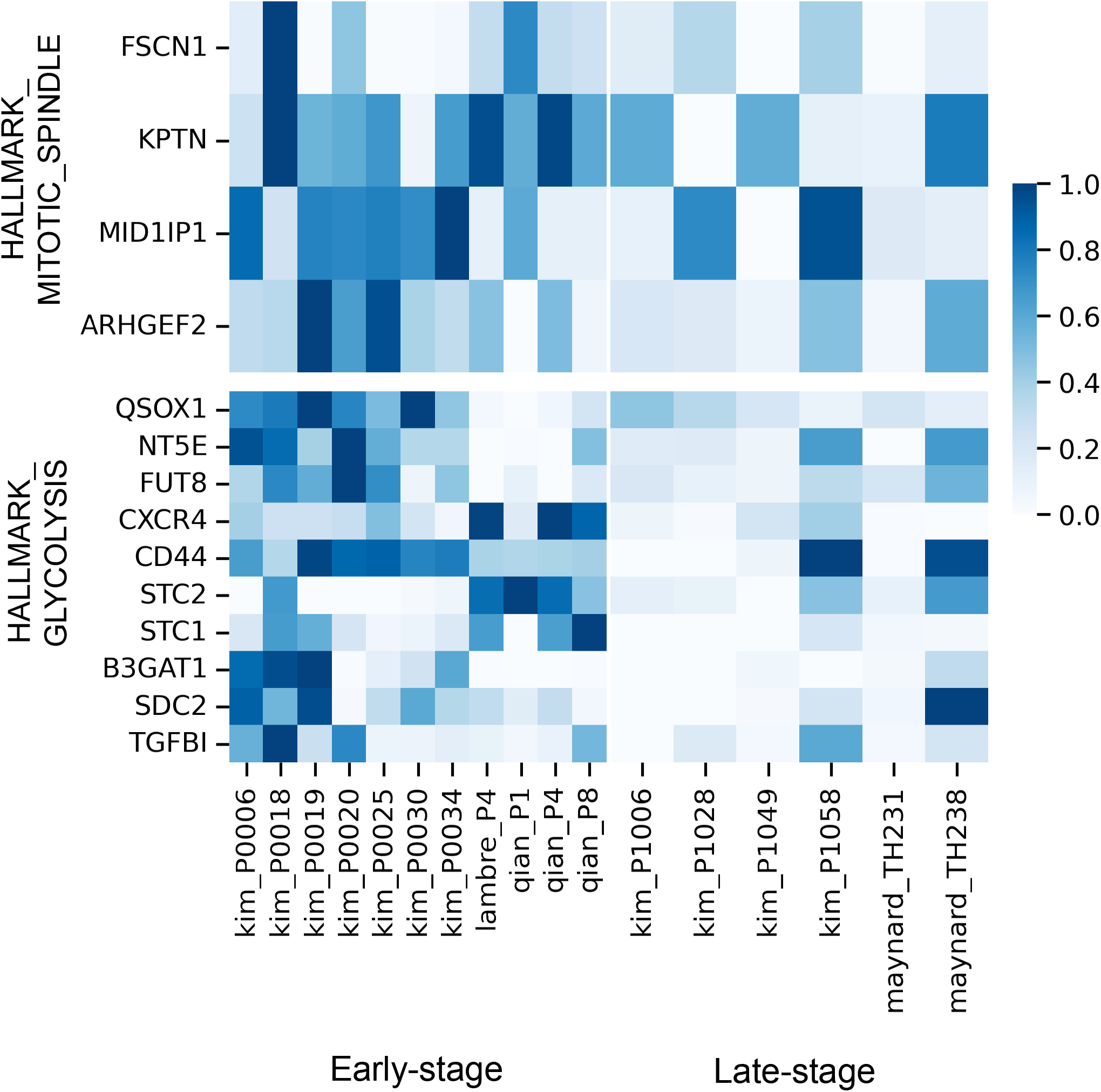
Heatmap displaying the differential top constituent genes (ranked according to log2 fold change and robustness) from the HALLMARK_MITOTIC_SPINDLE and HALLMARK_GLYCOLYSIS gene signatures in the early-vs. late-stage lung cancer dataset.

**Figure S3.**
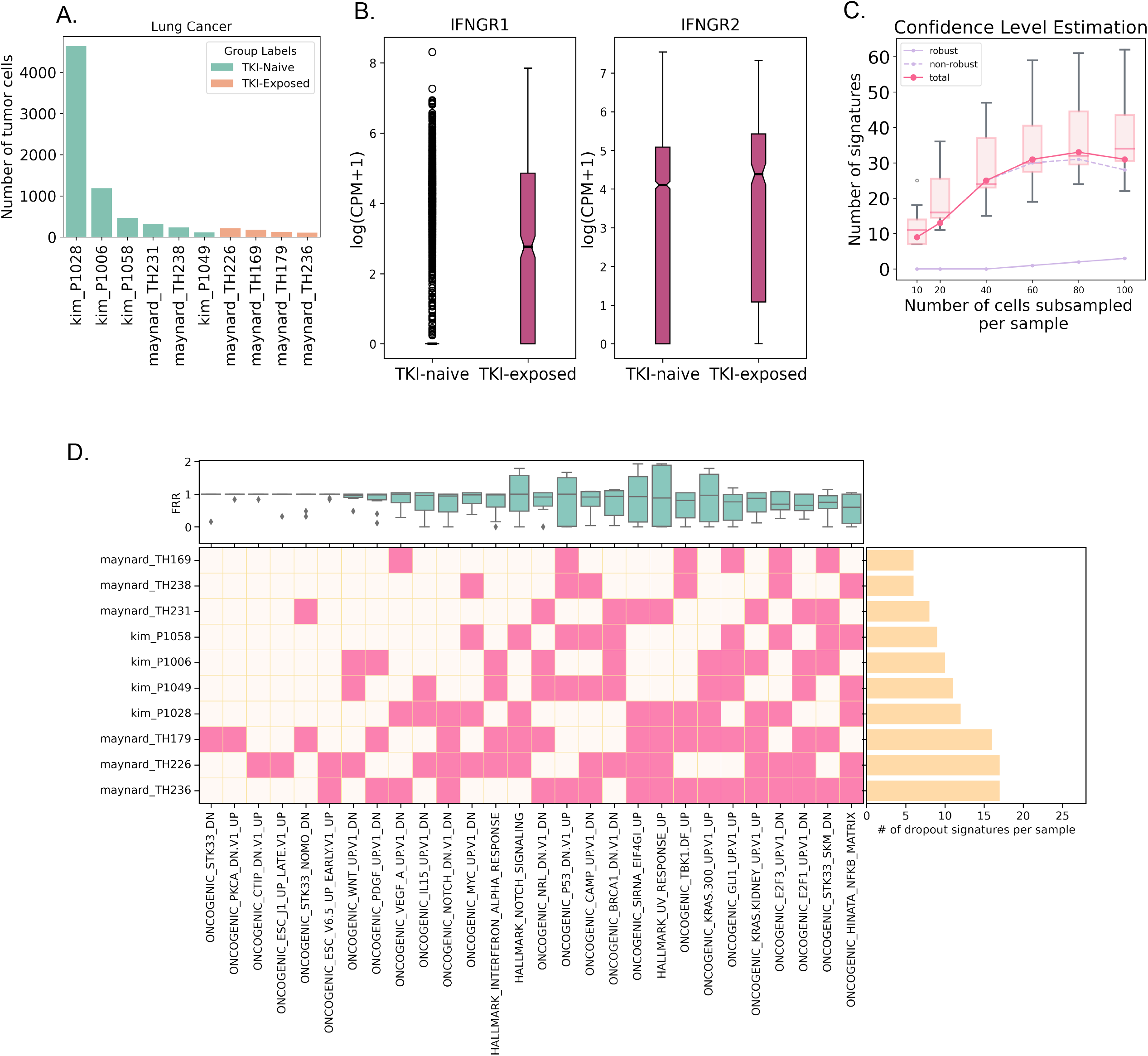
Extended group biology analysis of the tumor compartment from TKI-naive vs. TKI-exposed lung cancer patient samples. A. Distribution of tumor cells in the TKI-naive vs. -exposed lung cancer samples. B. Boxplot illustrating statistically significant differential expression of the genes IFNGR1 and IFNGR2 between TKI-naive vs. TKI-exposed samples. C. Line plot illustrating BEANIE’s stability to subsample size for TKI-naive vs. -exposed lung cancer samples. The curve plateaued as the number of signatures identified as statistically significant, irrespective of robustness status, reached saturation as the subsample size approached 100. D. Plot depicting the test signatures identified as statistically significant but non-robust to sample exclusion by BEANIE for TKI-naive vs. TKI-exposed samples, the distribution of their FRRs, and the patient IDs having FRRs less than threshold used to determine robustness, along with a horizontal bar plot of the number of statistically significant but non-robust signatures (dropout signatures) per sample.

## Supplementary Tables

Table S1: Hallmark and Oncogenic gene set results for MWU test + BH correction, GLMs, and BEANIE, for all datasets (ICB-naive vs. -exposed melanoma, early-vs. late-stage lung cancer, and TKI-naive vs. -exposed lung cancer).

Table S2: Top genes for Hallmark and Oncogenic gene sets for all datasets (ICB-naive vs. - exposed melanoma, early-vs. late-stage lung cancer, and TKI-naive vs. -exposed lung cancer).

Table S3: Noise estimation p-values for the three methods (MWU test with a BH correction, GLMs, and BEANIE).

Table S4: False positive rate (in percentage) for statistically significant and robust signatures identified by BEANIE for the Hallmark and Oncogenic gene sets for all datasets (ICB-naive vs. - exposed melanoma, early-vs. late-stage lung cancer, and TKI-naive vs. -exposed lung cancer).

## Notes

https://github.com/sjohri20/beanie

### Competing Interest Statement

The authors have declared no competing interest.

### Summary of Updates

Updated Abstract.

